# Association Mapping across Numerous Traits Reveals Patterns of Functional Variation in Maize

**DOI:** 10.1101/010207

**Authors:** Jason G Wallace, Peter J. Bradbury, Nengyi Zhang, Yves Gibon, Mark Stitt, Edward S. Buckler

## Abstract

Phenotypic variation in natural populations results from a combination of genetic effects, environmental effects, and gene-by-environment interactions. Despite the vast amount of genomic data becoming available, many pressing questions remain about the nature of genetic mutations that underlie functional variation. We present the results of combining genome-wide association analysis of 41 different phenotypes in ∼5,000 inbred maize lines to analyze patterns of high-resolution genetic association among of 28.9 million single-nucleotide polymorphisms (SNPs) and ∼800,000 copy-number variants (CNVs). We show that genie and intergenic regions have opposite patterns of enrichment, minor allele frequencies, and effect sizes, implying tradeoffs among the probability that a given polymorphism will have an effect, the detectable size of that effect, and its frequency in the population. We also find that genes tagged by GWAS are enriched for regulatory functions and are ∼50% more likely to have a paralog than expected by chance, indicating that gene regulation and neofunctionalization are strong drivers of phenotypic variation. These results will likely apply to many other organisms, especially ones with large and complex genomes like maize.

**Author Summary:** We performed genome-wide association mapping analysis in maize for over 40 different phenotypes in order to identify which types of variants are more likely to be important for controlling traits. We took advantage of a large mapping population (roughly 5000 recombinant inbred lines) and nearly 30 million segregating variants to identify ∼4800 variants that were significantly associated with at least one phenotype. While these variants are enriched in genes, most of them occur outside of genes, often in regions where regulatory variants likely lie. We also found a significant enrichment for paralogous (duplicated) genes, implying that functional divergence after gene duplication plays an important role in trait variation. Overall these analyses provide important insight into the unifying patterns of variation in traits across maize, and the results will likely also apply to other organisms with similarly large, complex genomes.

## Introduction

Natural phenotypic variation arises from a combination of genetic effects, environmental effects, and gene-by-environment interactions. A major goal of modem genetics is to tease apart these components, and especially to identify the genetic loci that govern variation in traits. In the past decade, genome-wide association studies (GWAS) have become a major tool to advance our understanding of genetic variation. While many genome-wide association studies (GWAS) focus on disease phenotypes, especially in humans (e.g., [1–5]), it is also important to identify the genetic nature of normal functional variation in populations-that is, all genetic variation which has a discernible phenotypic effect. There is also increasing evidence that differences in gene regulatory regions plays a significant role in functional variation [6–8], although the exact balance between regulatory variation versus protein-coding variation is still unsettled.

Because of the ability to create controlled crosses, model organisms provide powerful platforms to dissect this natural genetic variation. In recent years, large artificial populations have been created using several different organisms to leverage this power to dissect genetic traits (e.g., the mouse Collaborative Cross [9] and the Arabidopsis Multiparent Advanced Generation Intercross population [1OJ). Currently, the largest such population is the maize Nested Association Mapping (NAM) population [11]. Maize is an excellent genetic model for understanding natural variation due to the large phenotypic and genetic diversity available in its collections. NAM was designed to capture a large fraction of this variation by crossing 25 diverse founder lines to the reference line, B73, and generating 200 recombinant inbred lines (RILs) from each cross [11]. The hierarchical design of NAM provides both the high power of traditional linkage analysis and the high resolution of genome-wide association.

We leveraged the strengths of the NAM population to perform high-resolution GWAS across 41 diverse phenotypes to identify the general patterns of functional variation in maize. These traits were gathered from several individual studies on the NAM population (Table 1) and span the range of relatively simple metabolic traits up to highly complex traits such as height and flowering time. Our intent was not to re-identify regions influencing any specific trait, but rather to determine properties that make variants in general more likely to have a functional impact.

**Table 1:** Phenotypes used in this study

| *Phenotype* | *Citation* |
| --- | --- |
| Anthesis-silking interval | [14] |
| Average internode length (above ear) | [17] |
| Average internode length (below ear) | [17] |
| Average internode length (whole plant) | [17] |
| Boxcox-transformed leaf angle | [19] |
| Chlorophyll A | This study |
| Chlorophyll B | This study |
| Cob diameter | [12] |
| Days to anthesis | [14] |
| Days to silk | [14] |
| Ear height | [16] |
| Ear row number | [12] |
| Fructose | This study |
| Fumarate | This study |
| Glucose | This study |
| Glutamate | This study |
| Height above ear | [17] |
| Height per day (until flowering) | [17] |
| Kernel weight | Panzea.org<sup>a</sup> |
| Leaf length | [19] |
| Leaf width | [19] |
| Malate | This study |
| Nitrate | This study |
| Nodes above ear | [17] |
| Nodes per plant | [17] |
| Nodes to ear | [17] |
| Northern leaf blight | [18] |
| PCA of metabolites: PC1 | This study |
| PCA of metabolites: PC2 | This study |
| Photoperiod growing-degree days to anthesis | [15] |
| Photoperiod growing-degree days to silk | [15] |
| Plant height | [17] |
| Protein (total) | This study |
| Ratio of ear height to total height | [17] |
| Southern leaf blight | [13] |
| Stalk strength | [16] |
| Starch | This study |
| Sucrose | This study |
| Tassel branch number | [12] |
| Tassel length | [12] |
| Total amino acids | This study |

<sup>a</sup> http://www.panzea.org/lit/data_sets.html#phenos ; the joint-linkage model to create residuals
for this data was provided courtesy of Sherry Flint-Garcia

We expect to have very high resolution for these hits because of the speed with which linkage disequilibrium (LD) decays in maize. An empirical calculation of LD decay in NAM shows that most LD decays to below background levels within 1 kilobase of a given polymorphism, though the variance is large since some alleles are segregating in only one or two families (Supplementary Figure 1). Due to this rapid LD decay, the high density of polymorphisms we used, and the high statistical power gained by using the NAM population, we expect that many of the polymorphisms we identified will be extremely close (within a few kb) to the causal polymorphism. Furthermore, since our polymorphism dataset covers much of the low-copy fraction of the genome, some unknown fraction of these hits will probably be the causal polymorphisms themselves (though we cannot currently tell which ones they are).

We find that a large amount of functional variation is located outside of protein-coding genes, presumably in regulatory regions, and that these non-genie variants often have large phenotypic effects. We also find that genes identified by association analysis are enriched for regulatory functions and for paralogs; this latter implies that neofunctionalization (acquiring a novel function after gene duplication) is likely to be a strong driver of normal functional variation.

## Results

### Phenotype data

The majority of phenotype data in this analysis was taken from existing studies on the maize Nested Association Mapping population (Table 1) [12–19]. These existing phenotypes cover various plant architecture, developmental, and disease resistance traits. In addition, we also obtained trait data for 12 different metabolites in leaves: Chlorophyll A, Chlorophyll B, Fructose, Fumarate, Glucose, Glutamate, Malate, Nitrate, Starch, Sucrose, Total amino acids, and Total protein. (Details of data acquisition are in the Methods section.) An in-depth analysis of these metabolites and the variants associated with each of them is forthcoming (Zhang *et al.,* in preparation); for this paper, we used them primarily to expand our pool of available phenotypes. Both raw metabolite data and best linear unbiased predictors (BLUPs) for each NAM line are included in Supplemental File 1.

### Genome-wide association

Single-nucleotide polymorphisms (SNPs, also including short indels of <15 base pairs) were taken from Maize Hapmapl [20] and Hapmap2 [21], for a total of 28.9 million segregating SNPs. We also used the raw Hapmap2 read depth counts to identify ∼800,000 putative copy number variants (CNVs) as done previously [21].

These 29.7 million total segregating polymorphisms were then projected onto the 5,000 RIL progeny based on low-density markers obtained through genotyping-by-sequencing (GBS) [22]. We then performed forward-regression GWAS to identify which of these variants associated with the different phenotypes. Full details are in the Methods section; in brief, the forward-regression model iteratively scans the genome, each time adding only the most significant SNP to the model until no SNPs pass the significance threshold. We ran 100 such genome-wide associations for each trait with a random 80% of lines subsampled each time. The random subsampling allows us to filter based on how many iterations a SNP appears in, a measure of the strength and stability of the association.

After filtering to remove hits that showed up in <5 iterations [12,13], we identified 4,484 SNPs and 318 CNVs that were significantly associated with at least one phenotype. The number of polymorphisms identified for each trait varies widely and matches prior assumptions based on the genetic complexity of the traits (Figure 1). Comparing our results with those of published studies in NAM shows good agreement with the locations of known QTL (Supplementary Figure 2).

**Figure 1:**
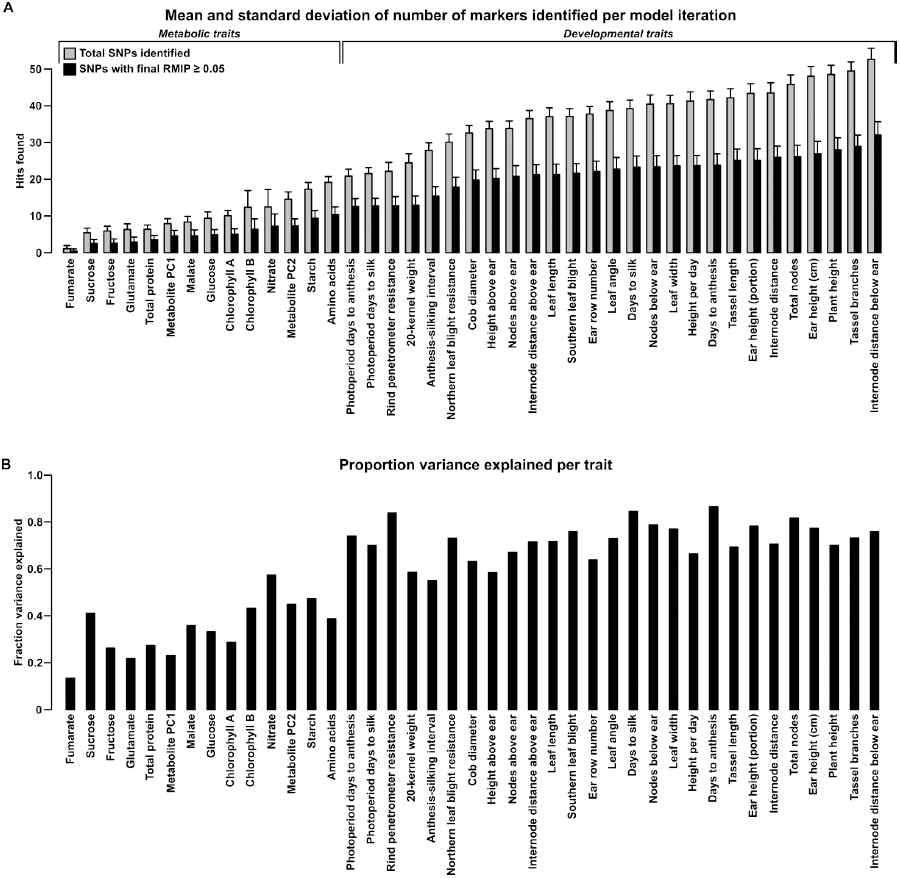
Number of polymorphisms found and variance explained for each trait. (A) Polymorphisms found per trait. Bars show the mean and standard deviation of markers found per iteration before (light bars) and after (dark bars) filtering for RMIP≥ 0.05 (see Methods). The number of markers found tends to mirror the genetic complexity of each trait, with metabolic traits having fewer markers found than complex, polygenic traits like plant architecture. (B) Variance explained per trait. For each trait, a general linear model incorporating a family term (for each of the 25 biparental families in NAM) and all SNPs that passed filtering (dark bars in (A)) was fit to the original Best Linear Unbiased Predictors (BLUPs) for each trait. Bars show the portion of total variance explained by the fitted SNPs as measured by adjusted R^2^

### Variant classification

To classify each polymorphism, we used the Ensembl Variant Effect Predictor (VEP) [23] to identify the potential effect of each SNP in both the input and GWAS datasets. Since most SNPs are likely not causal but just linked to the causal polymorphism, these annotations serve primarily to identify the region a SNP lies in and the types of SNPs most frequently identified by GWAS across our dataset.

After classification, we analyzed the distribution of VEP classes and copy-number variants (CNVs) for enrichment in GWAS hits relative to the input dataset (Figure 2). Intergenic regions (>5 kb away from the nearest gene) are strongly depleted for GWAS hits, causing almost all other categories to show significant enrichment (Figure 2B). Part of this depletion may be due to transposon activity in intergenic regions altering the physical location-and thus the projected genotype—of sequences in some founder lines. After controlling for intergenic regions, both genie SNPs and CNVs are still strongly enriched for GWAS hits (Figure 2C). This agrees with the recent findings of Schork *et al.* (2013), who found similar enrichment patterns of GWAS hits close to genes. Of the enriched classes, large CNVs show the most enrichment, while the most enriched SNP category is for synonymous mutations. Some of the enrichment for synonymous sites is probably due to synthetic associations [24], where a high-frequency, synonymous SNP is identified instead of several nearby, low-frequency mutations. However, synonymous SNPs are also significantly enriched over intronic SNPs (p=2.80x10^-8^ by Chi-square test) despite having similar site frequency spectra (data not shown) and being in similar LD structures (due to the small size of maize introns, which have a median size of only ∼150 base pairs in quality-filtered genes). This implies a legitimate enrichment for synonymous SNPs. Some (and possibly most) of that enrichment is probably due to linkage with nearby causal SNPs, while the remainder is likely due to the (unknown) fraction that are causal themselves but act through mechanisms other than protein sequence (e.g., altering mRNA stability, protein binding sites, or local translation rates [25]).

**Figure 2:**
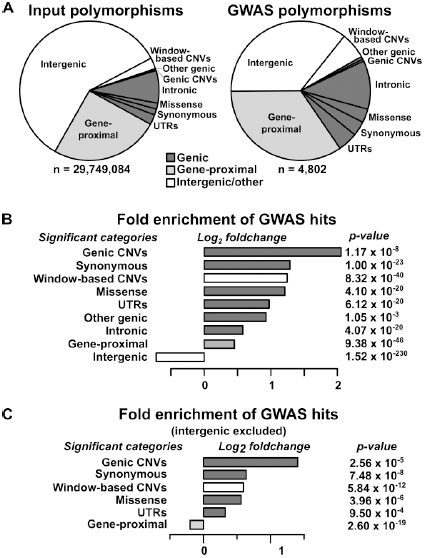
Relative enrichment of polymorphism classes in GWAS hits. (A) The proportions of different polymorphism classes in the input dataset (left) and GWAS hits (right). The overall GWAS hit distribution is significantly different from the input at p = 8.74x10·^35^ (Chi-square test). (B) The relative change in polymorphism classes in the GWAS dataset as compared to the input dataset, with the raw p-value of each class shown at right (two-sided exact binomial test). Only categories with Bonferroni-corrected p-values ≤ 0.01 are shown. The strong depletion of intergenic SNPs in the GWAS dataset drives almost all other categories to appear significantly emiched. Exact category counts and alternate p-values based on circular permutation are available in Supplementary Table 1. (C) The same analysis as in (B), but with intergenic regions excluded.

Although genie regions are the most strongly enriched in GWAS, the majority (∼70%) of our hits still fall outside of annotated genes, as defined by their transcriptional start and stop sites. Plotting the distances from non-genie SNPs to the nearest gene on a log scale reveals a bimodal distribution, with a peak at ∼1-5 kb away from genes that is not reflected in the input dataset (Figure 3). This corresponds with likely positions of promoters and other short-range regulatory elements. Finding enrichment at this scale provides evidence for the high resolution and biological relevance of the GWAS hits in this study. The second peak, which follows the null distribution, probably reflects elements that are not correlated with gene distance (e.g., long range regulatory elements, unannotated transcripts, etc.). For example, using a list of 316 maize noncoding RNAs from Gramene (available at http://ftp.gramene.org/release39/data/fasta/zea_mays/ncma/) that were not included in the Ensembl annotations reveals that intergenic hits are significantly enriched for polymorphisms within 5 kb of these RNAs (n=13, expected=l.07, p=l. 3x10^-10^ by two-sided exact binomial test). Alternatively, some of these “intergenic” hits may actually be tagging legitimate genes that are simply not present in the reference genome due to the high amount of presence-absence variation in maize [21]. Identifying the nature of these hits should be possible as more information about the maize pan-genome becomes available.

**Figure 3:**
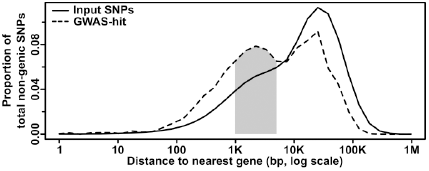
Distribution of non-genie GWAS hits as a function of gene distance. The number of SNPs at increasing distances from the nearest gene is plotted; CNVs are excluded due to their large size and the difficulty determining where many (especially insertions) actually occur. The input (whole genome) dataset shows a single peak at ∼25 kb away from a gene. The GWAS dataset, however, shows an additional peak at ∼1-5 kb (shaded), where one would expect to find promoters and short-range regulatory elements. Note that due to the log scale, each bin contains successively more nucleotides that make it appear that most SNPs are far from genes, when the reverse is actually true.

### Relative effect sizes of the different classes

We also determined the relative effect each polymorphism class has on phenotype. We classified all SNP hits by whether they fell within genes (genie), within 5 kb of a gene (gene-proximal), or more than 5 kb away (intergenic), and compared the variance explained among traits for these classes and for CNVs (Figure 4A). Genie and gene-proximal SNPs explain the most unique variance, meaning the proportion of variance explained when the specified category is added last to a model. However, examining the minor allele frequency (MAF) and effect size distributions for each class reveals a more complex picture (Figures 4B & 4C). Both MAF and effect size strongly influence variance explained, and in our dataset they are negatively correlated. Similar results were found in a previous study of inflorescence traits [12]. This negative correlation is probably due to both biological factors (e.g., large-effect mutations are more likely to be detrimental to overall fitness [26,27] and thus kept at low frequency) and also statistical limitations (e.g., GWAS can only identify rare variants if they have large effects). At the extremes, intergenic variants have the largest median effect size but the lowest allele frequencies, while CNVs are the reverse. Thus many large phenotypic effects tend to occur outside of genes (presumably in regulatory elements, unannotated transcripts, or the like), but they also tend to be rare and so make only minor contributions to total variance explained. This inverse relationship between allele frequency and effect size holds across polymorphism classes (Figure 5), implying a general pattern across polymorphisms. Since large-effect polymorphisms are exactly the sort of mutation breeders often look for in selecting germplasm for breeding programs, these data may prove useful for future breeding efforts.

**Figure 4:**
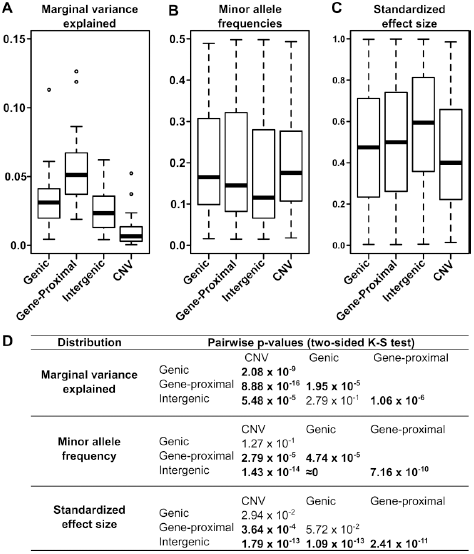
Figure 4: Different effects of the polymorphism classes. (A) Variance explained by polymorphism class. Genie and gene-proximal polymorphisms explain the largest amount of unique variation in each trait. Breaking the data into the two components that most influence variance explained-allele frequency (B) and polymorphism effect size (C)-reveals a negative correlation between them such that classes with larger effect sizes (e.g., intergenic) also tend to have rarer polymorphisms. (D) Pairwise p-values testing whether the distributions in (A-C) are significantly different from each other (two-sided Kolmogorov-Smirnov test); values< 1 x 10−^3^ are bolded.

**Figure 5:**
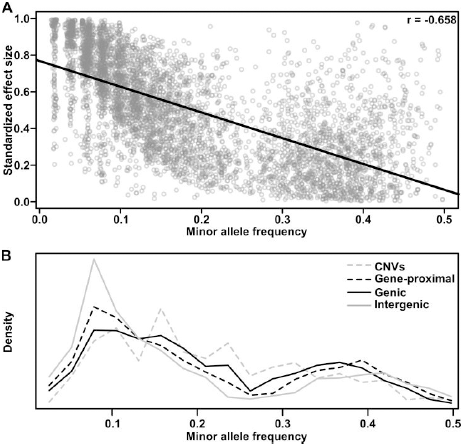
Polymorphism effect size and allele frequencies. (A) The standardized effect size of a polymorphism (see Methods) is negatively correlated with minor allele frequency. This correlation is probably due to both biological factors (e.g., large effects are both more likely to deleterious (Fisher 1930; Orr 1998) and more easily selected against than small ones, and thus are more likely to remain rare) and statistical ones (e.g., in order for a rare variant to explain enough variance to be detected in GWAS, it must have a large effect). Similar results were found in a previous analysis of maize inflorescence traits [12]. (B) Minor allele frequency distributions for the different polymorphism classes of GWAS hits. Intergenic hits are strongly enriched for rare alleles. The bimodal distribution in both parts is due to the way NAM was constructed; specifically, since B73 is a parent in all 25 families, any polymorphisms with the rare allele in B73 have their frequency artificially boosted toward 0.5.

### Characteristics of GWAS-hit genes

Since the annotation of single nucleotides in genie regions is more straightforward than in intergenic regions, we also identified common characteristics of genes that were tagged by genie or gene-proximal GWAS hits.

First, an analysis of expression levels using RNA-seq data from the Maize Gene Atlas [28] reveals a small (∼20%) but highly significant depletion oflow-expressed genes (p=l.30x10^-22^ by Mann-Whitney test and ≈ 0 by Kolmogorov-Smirnov test) (Figure 6). The expression level of these genes is even lower than most transcription factors, which are themselves usually only expressed at a low level, and their depletion among GWAS hits may reflect a lower probability of such rarely expressed genes altering plant phenotype.

**Figure 6:**
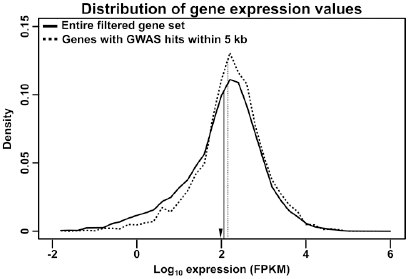
Distribution of RNA expression. Transcript-specific RNA expression values from the Maize Gene Atlas [28] were summed to determine total expression for each gene. The log transformed distribution of maximum expression values are shown for the entire filtered gene set (solid line) or just genes with GWAS hits within 5 kb of their primary transcripts (dashed line); vertical lines indicate the median of each distribution. The GWAS-hit genes show a slight depletion (∼20%) of low-expressed genes. For comparison, the median expression of maize transcription factors in this dataset (as annotated on Grassius, http://grassius.org/) is indicated by an arrowhead. FPKM, Fragments Per Kilobase of transcript per Million mapped reads.

Second, Gene Ontology (GO) term analysis revealed significant emichment (∼34%) in terms relating to regulatory activity, especially protein kinase activity and transcription factor activity, and depletion (∼71%) among several core metabolism and signaling terms (Supplementary Table 2). These terms are fairly broad, probably because the diverse phenotypes in this study make it so that the only terms that are significantly changed are those general enough to be involved across many different phenotypes. Nonetheless, the enrichment of regulatory terms across such a broad phenotypic spectrum implies that changes in gene regulation are a frequent driver of functional variation. Conversely, the depletion of core metabolic terms speaks to the difficulty of altering these functions without causing detriment to the organism.The depletion in core metabolic terms is especially striking because the studied traits include 12 metabolic traits.

Finally, we found that genes with GWAS hits in their primary transcripts are ∼50% more likely to have a paralog than expected by chance (36.4% of 970 GWAS-hit genes vs 24.2% of 39,656 total genes in the filtered gene set; p=3.79x10^-17^ by two-sided exact binomial test). Paralogous genes do not appear to have significant differences from non-paralogous genes in either allele frequency or LD structure, and the marginally lower density of SNPs in them would seem to disfavor their selection by GWAS, all other things being equal (Figure 7). Thus the enrichment for paralogous genes is probably due to the benefits of neofunctionalization, where having redundant copies of a gene allows one of them to more easily take on altered (and phenotypically significant) roles [29].

**Figure 7:**
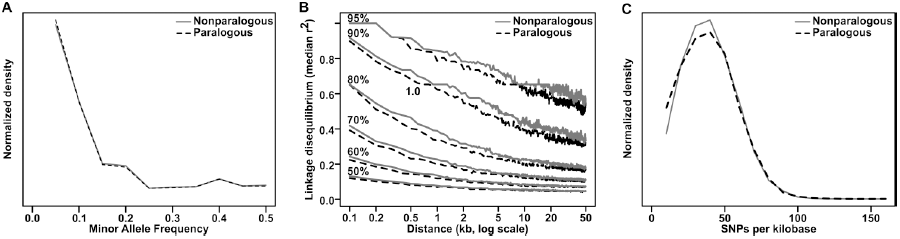
Comparison of paralogous to nonparalogous genes. Maize paralogous genes (identified by Schnable & Freeling [50]) were examined for any differences from nonparalogous genes that might spuriously contribute to their emichment in GWAS analyses. There are no strong differences in either minor allele frequency distribution (A) or linkage disequilibrium decay (B), and the slightly lower SNP density (C) (median 32.8 SNPs/kb versus 33.4 SNPs/kb for nonparalogous genes) would be expected to actually decrease the probability of hitting paralogous genes, albeit by a very small amount.

## Discussion

Taken together, the large number and effect sizes of hits outside genes and the enrichment for copy-number variants indicate that while variation in gene sequence is important, a large portion of functional variation in maize probably stems from differences in copy number and gene regulation rather than in protein-coding sequence. These results corroborate similar findings in other organisms [6–8], indicating that this pattern will likely hold for many other species. One caveat, however, is that our filtering for robust GWAS hits intrinsically skews the results toward more common alleles; rare variants may follow different patterns.

Our results also imply that the cost-saving measure of genotyping individuals by sequencing only the exome may be of limited utility for GWAS, at least for organisms like maize where LD decays rapidly. This is in direct contrast with the conclusions of Li *et al.* [30], who determined that 79% of the explained variation in their maize dataset could be encompassed by genie and promoter (<5 kb upstream) regions. We suspect that this difference is chiefly due to choice of input polymorphisms*. Li et al.* used-290,000 SNPs derived from RNA-seq data and <775,000 SNPs from Maize Hapmapl; the former is obviously biased toward genie regions, while the latter has a similar (albeit smaller) bias due to using methyl-sensitive restriction enzymes to construct genomic libraries [20]. In contrast, the majority (∼92%) of our input polymorphisms come from Maize Hapmap2, where sequencing libraries were created by random shearing and thus show much smaller bias toward genie regions [21].

Ultimately, the goal of modem crop genetics is to design crops for rapidly changing environments. Doing so requires accurate information about which genomic regions contribute to trait qualities. The fact that most of our hits (70%) lie in poorly annotated regions outside of annotated genes and that these hits often have large phenotypic effects argues for an urgent need to identify the genetic features in these regions. Such efforts are already underway for humans and several model animals [31–33]; similar work should be extended to plants and especially to important crops like maize. The low cost of current sequencing would even make it possible to, for example, combine GWAS with expression profiling across several thousand individuals to identify both regulatory regions and their effects on phenotype. Identifying these features and including them in prediction models will further not only basic genetics, but also help breeders craft better crops and help improve food security for the global population.

## Methods

### Bioinformatics and statistics

Unless otherwise stated, all analyses were performed with in-house bioinformatics pipelines written in SAS, R, Perl, or Java. Source code is available upon request. All analyses were done with using the maize B73 genome (version AGPv2) as reference.

### Metabolite data

#### Sampling

The NAM population was planted in Aurora, New York, USA in May 2007. Samples were all taken within one week at the beginning of August (when most NAM lines are flowering) between 10:00 AM and 2:00 PM on the sampling date. Two samples were taken from each row (RIL), one from the end plant and the other from four middle plants (∼12,000 samples total). Tissue was punched in the base part of the first leaf below the flag leaf and immediately frozen in liquid nitrogen, then stored at -80°C until extraction.

#### Quantification

∼50 mg (fresh weight) of tissue was extracted twice with 80% ethanol and once with 50% ethanol as in Geigenberger et al. [34] (the final volume of each was 650 µl). Protein and starch were extracted from the pellet with 100 mM NaOH [35] and measured according to established protocols [35,36]. Immediately after extraction, chlorophyll content was determined using the protocol in Amon [37]. Total free amino acids were assayed using fluorescamine [38]. Nitrate levels were quantified as in Tschoep *et al.* [39], *while malate and fumarate were measured as described in Nunes-Nesi et al.[40].* Glutamate was determined by pipetting 10 µl aliquots of extract or standards (0-20 nmol) into a microplate with 100 mM Tricine/KOH pH 9, 3 mM NAD+, lmM methylthiazolyldiphenyl-tetrazolium bromide, 0.4 mM phenazine ethosulphate and 0.5% v/v Triton X-100. The absorbance at 570 nm was read for 5 min, then 1 U of glutamate dehydrogenase was added and the absorbance monitored until it reached stability. Sucrose, glucose, and fructose (in ethanolic extracts) were determined as per Jelitto *et al.* [41]. All assays were prepared in 96-well polystyrene microplates using a JANUS automated workstation robot (Perkin-Elmer, Zaventem, Belgium). Absorbances at 340 or 570 nm were read in either an ELX-800 or an ELX-808 microplate reader (Bio-Tek, Bad Friedrichshall, Germany). A Synergy microplate reader (Bio-Tek, Bad Friedrichshall, Germany) was used to determine absorbances at 595, 645 or 665 nm and fluorescence (405 nm excitation, 485 nm emission).

#### BLUPs and Principal Components

Best linear unbiased predictors (BLUPs) for each line within each trait were calculated using ASReml (version 2.0; http://www.vsni.co.uk/software/asreml). The final BLUPs are the result from controlling for several potential confounding factors, specifically: spatial effects within the field; the level of nitrogen, phosphorous and potassium in the soil before planting; the tissue sampling date and time; the researcher who performed the sampling; the batch effect of the plate samples were stored in; and the batch effect of the plate the measurements occurred in. BLUPs were also calculated for flowering time (defined as the time from sowing to when 50% of plants in a row are shedding pollen), correcting for the spatial field effects. Most metabolites correlate with flowering time (data not shown), so we performed partial correlation analyses with Proc GLM in SAS (http://www.sas.com/) to account for its effect on the 12 metabolites. A sequential Bonferroni test [42] at α = 0.05 was used to correct for multiple testing. Principal components were calculated with Proc PRINCOMP in SAS after fitting a linear model to account for the effect of flowering time (days to anthesis) as a covariate. (That is, the principal components are of the residuals after factoring out flowering time.)

### GWAS Analysis

Phenotype data for GWAS analysis was taken from previous studies by our lab and others on a variety of traits, along with the metabolite data included herein (Table 1). In the majority of cases phenotypic data had already been processed by fitting a joint-linkage model [43] with 1,106 high-confidence SNP markers across NAM. Chromosome-specific residuals were then determined by fitting a model that included as covariates all identified quantitative trait loci (QTL) except those on the given chromosome. For traits without precomputed residuals, the same process was followed but with an updated list of ∼7,000 SNPs derived from genotyping by-sequencing [22]. All genotypes are available at http://www.panzea.org; chromosome-specific residuals are included in Supplemental File 2.

Forward-regression genome-wide association was then performed with the NamGwasPlugin in TASSEL version 4.1.32 [44]. Each chromosome was analyzed separately for each phenotype via 100 forward-regression iterations, each of which excluded a random 20% of NAM lines to destabilize spurious associations [45]. The cutoff for polymorphism inclusion in the model was a raw p-value <9.50x10^-8^, which was empirically determined by permutation testing with the days to anthesis phenotype to correspond to a genome-wide Type I error rate of 0.01. The resample model inclusion probability (RMIP) [45] of each polymorphism was determined as the proportion of iterations in which a specific polymorphism was called as significant; only polymorphisms with an RMIP e 0.05 are considered in this study. We found a single case of ambiguity in determining which SNP had been chosen by the model, due to two SNPs having the same position and allele codings but different original sources (Hapmap1 vs Hapmap2). In this case we retained both to maintain consistency with the input dataset.

### Copy-number variants

Putative CNVs were determined by two methods. First, Hapmap2 sequencing reads aligned to the maize genome were counted in 2 kb-windows and compared to a high-coverage B73 sample with edgeR [46]. This procedure had been done previously [21], and our analysis was primarily to update the results to a newer version of the *Zea mays* reference genome (AGPv2). The B73 sample from Hapmap2 itself served as the null distribution to determine the cutoff corresponding to an empirical, genome-wide Type I error rate of 0.05. CNVs that had been previously identified within annotated genes by the same method [21] were also included in the analysis.

Independently, the mapped reads were also analyzed by CNVnator [47] to identify putative CNVs based on shifts in mean read depth across 500 bp bins. Interestingly, although many CNVnator CNVs showed consistent segregation across the NAM founders, GWAS hits came almost exclusively from the edgeR-derived CNVs. Looking at the characteristics of each, this disparity is probably due to two factors: (1) the edgeR-derived CNVs are generally much smaller than those found by CNVnator, and smaller CNVs have previously been shown to have more significant GWAS hits in this population [21]; and (2) edgeR also detects many more CNVs than CNVnator to begin with, presumably because small CNVs are more common than large ones.

### SNP annotation

Putative SNP effects were determined by running all SNPs through the Ensembl Variant Effect Predictor (VEP) [23] using a local copy of the *Zea mays* Ensembl database (version 68). Since the VEP annotates effects relative to any gene model (not just quality-filtered ones), it was run with both the “`--most-severe`” and “`--per-gene`” options to get lists of the worst overall effect per SNP and the worst per gene, respectively. (Note that the VEP considers that changing an existing amino acid is more severe than in-frame insertions and deletions, so small indels that do both get classified as “missense.” These make up <0.1% of the input polymorphisms and only 3 GWAS hit ones, however, so altering the annotation would not significantly affect the results.) The two results were then combined with in-house Perl scripts to create a list of the worst overall SNP effect with respect to only those genes in the *Zea mays* 5b.60 filtered gene set (available at http://www.gramene.org).

### Polymorphism class enrichment

Using the input dataset as the null distribution, the overall significance of the difference in category distributions was determined by a Chi-square test using the Stats package in R [48]. Individual categories were then tested for emichment by a two-sided exact binomial test, also in R.

Due to the possibility that linkage disequilibrium could distort the results from the above test, we also ran 1 million circular permutations of the hits to generate a null distribution of what would be expected by chance. The resulting counts formed a normal distribution, which was used to extrapolate the p-values in Supplementary Table 1.

### Marginal variance explained

Marginal variance explained by polymorphisms classes (genie, gene-proximal, intergenic, and CNVs) was calculated by fitting linear models to each trait and comparing the difference in variance explained (adjusted R^2^) between a model with all identified SNPs and a model with all SNPs except those in the chosen category.

### Standardized effect sizes

Standardized effect sizes for each polymorphism were determined by first taking all effect sizes the NAM-GWAS model identified for each trait and fitting an empirical cumulative distribution function with ecdf() in R [48]. This function was then used to determine the quantile of each effect. Mean quantile scores were then calculated for each polymorphism that passed RMIP ≤ 0.05 filtering. Each point in the distribution thus represents a specific trait-polymorphism combination.

### GO *Term Enrichment*

Gene Ontology term analysis was performed with agriGO [49] using all genes with GWAS hits within 5 kb of their annotated transcript. Statistical analysis was performed in R [48] via a two sided Fisher’s exact test with Benjamini-Yekutieli control of the false discovery rate (FDR) to analyze for both emichment and depletion.

### Paralogy

Maize paralogs were taken from an existing list [50] (available at http://genomevolution.org/CoGe). The number of genes with paralogs in the GWAS hit dataset was compared to those in the maize filtered gene set and significance of the difference tested by a two-sided exact binomial test in R [48].

## Acknowledgements

We thank the participants of the Panzea project (http://www.panzea.org) for gathering the phenotypic and genotypic data used in this study; Jason Peiffer and Sherry Flint-Garcia for sharing prepublication data; Jer-Ming Chia for the raw Hapmap2-aligned reads; and Sara Miller for assistance in editing the manuscript.

## Supplemental Data

**Figure S1:**
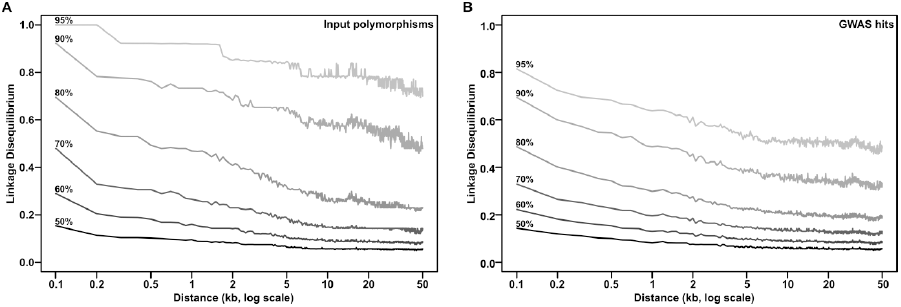
Linkage disequilibrium in NAM. Linkage disequilibrium (LD) in the NAM population was calculated for 10,000 random polymorphisms (A) and for all GWAS hits (B) based on expected contribution from the 26 founder genotypes. Lines show the distribution of polymorphisms at different percentile cutoffs (marked at left). Median LD, as marked by the 50% line, falls below background (r^2^ < 0.2) in less than 100 base pairs. Rare variants segregating in just a few lines create a large variance in LD structure, however, as shown by the persistence of LD at higher percentile cutoffs.

**Figure S2:**
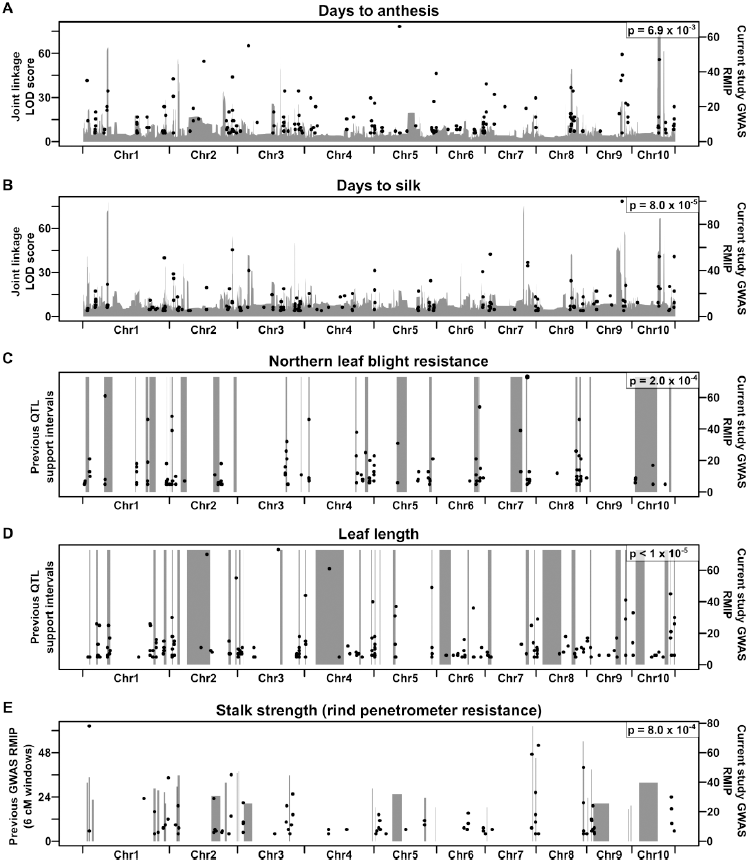
Agreement between identified polymorphisms and known QTL. Quantitative trait loci (QTL) for key traits from previous studies in NAM were compared against polymorphisms found in the current analysis (black dots). Gray bars show the results of genome wide joint-linkage scans for days to anthesis (A) and days to silk (B) (Buckler *et al.* 2009), QTL support intervals for Northern leaf blight resistance (C) and leaflength (D) (Poland *et al.* 2011; Tian *et al.* 2011), and 6 cM windows of significant SNPs for stalk strength (E) (Peiffer *et al.* 2013). 100,000 circular permutations were performed to determine the significance of overlap between the previous results and our GWAS hits; the resulting empirical p-values are in the upper-right of each graph. (Since parts (A) and (B) are continuous scans, a LOD-score cutoff of 15 was used to specify QTL intervals.) All overlaps are significant at p<0.01. It should be noted that the lack of perfect overlap is largely due to the different statistical strengths of joint linkage and GWAS, and similar results are seen in the previous NAM studies that used both methods.

**Table S1:** Category counts

| Category | Input counts | GWAS counts | Permutation counts<sup>ab</sup> | Enrichment<sup>a</sup> | p-value<sup>a</sup> |
| --- | --- | --- | --- | --- | --- |
| Gene-proximal SNPs | 7,505,673 | 1,654 | 1,166.1 ± 75.56 | 1.42 | 1.07 x 10<sup>-10</sup> |
| Genic CNVs | 35,743 | 24 | — | — | — |
| Intergenic SNPs | 17,570,581 | 1,720 | 2,714.8 ± 118.95 | 0.63 | 6.11 x 10<sup>-17</sup> |
| Intronic SNPs | 2,246,611 | 542 | 350.5 ± 30.98 | 1.55 | 6.38 x 10<sup>-10</sup> |
| Missense SNPs | 409,728 | 153 | 63.7 ± 9.46 | 2.4 | 3.85 x 10<sup>-21</sup> |
| Other genic SNPs | 100,940 | 31 | 15.7 ± 4.18 | 1.98 | 2.47 x 10<sup>-4</sup> |
| Synonymous SNPs | 416,452 | 164 | 64.9 ± 9.77 | 2.53 | 3.62 x 10<sup>-24</sup> |
| UTR SNPs | 694,926 | 220 | 108.3 ± 14.40 | 2.03 | 8.61 x 10<sup>-15</sup> |
| Window-based CNVs | 768,430 | 294 | — | — | — |
| *total* | 29,749,084 | 4,802 |  |  |  |

<sup>a</sup>One million circular permutations of each chromosome were performed to determine empirical enrichment of each SNP category; CNVs were excluded due to ambiguity in their precise placement on chromosomes, especially for duplications. The resulting normal distribution of counts was used to extrapolate two-sided p-values for enrichment, since the actual values were generally more extreme than any observed permutation.
<sup>b</sup>mean ± standard deviation among all permutations

**Table S2:** GO term analysis

| GO term | Description | GWAS hits<br>(n=1,879 total terms) | Whole-genome hits<br>(n=25,288 total terms) | FDR<sup>a</sup> | Odds ratio |
| --- | --- | --- | --- | --- | --- |
| **Enriched terms** |  |  |  |  |  |
| *Protein kinase-related* |  |  |  |  |  |
| GO:0004713 | Protein tyrosine kinase activity | 150 | 1313 | 7.58x10<sup>-05</sup> | 1.66 |
| GO:0004674 | Protein serine/threonine kinase activity | 160 | 1430 | 8.92x10<sup>-05</sup> | 1.62 |
| GO:0006468 | Protein amino acid phosphorylation | 167 | 1528 | 1.89x10<sup>-04</sup> | 1.58 |
| GO:0004672 | Protein kinase activity | 168 | 1552 | 3.38x10<sup>-04</sup> | 1.56 |
| GO:0016301 | Kinase activity | 195 | 1909 | 1.87x10<sup>-03</sup> | 1.47 |
| GO:0016773 | Phosphotransferase activity, alcohol group as acceptor | 186 | 1824 | 2.59x10<sup>-03</sup> | 1.46 |
| GO:0043687 | Post-translational protein modification | 186 | 1832 | 3.13x10<sup>-03</sup> | 1.45 |
| GO:0043412 | Macromolecule modification | 204 | 2069 | 7.16x10<sup>-03</sup> | 1.41 |
| GO:0006464 | Protein modification process | 195 | 2013 | 2.96x10<sup>-02</sup> | 1.38 |
| GO:0032559 | Adenyl ribonucleotide binding | 292 | 3197 | 3.04x10<sup>-02</sup> | 1.30 |
| GO:0005524 | ATP binding | 292 | 3193 | 3.04x10<sup>-02</sup> | 1.30 |
| GO:0016310 | Phosphorylation | 178 | 1821 | 3.10x10<sup>-02</sup> | 1.39 |
| GO:0001883 | Purine nucleoside binding | 304 | 3384 | 4.83x10<sup>-02</sup> | 1.27 |
| GO:0001882 | Nucleoside binding | 304 | 3385 | 4.83x10<sup>-02</sup> | 1.27 |
| GO:0030554 | Adenyl nucleotide binding | 304 | 3384 | 4.83x10<sup>-02</sup> | 1.27 |
| *Transcription factor-related* |  |  |  |  |  |
| GO:0030528 | Transcription regulator activity | 153 | 1432 | 1.87x10<sup>-03</sup> | 1.53 |
| GO:0003700 | Transcription factor activity | 106 | 923 | 2.37x10<sup>-03</sup> | 1.65 |
| GO:0010468 | Regulation of gene expression | 210 | 2216 | 3.81x10<sup>-02</sup> | 1.34 |
| GO:0043565 | Sequence-specific DNA binding | 74 | 645 | 3.81x10<sup>-02</sup> | 1.64 |
| GO:0045449 | Regulation of transcription | 205 | 2172 | 4.83x10<sup>-02</sup> | 1.33 |
| *General regulation* |  |  |  |  |  |
| GO:0009889 | Regulation of biosynthetic process | 209 | 2210 | 3.90x10<sup>-02</sup> | 1.34 |
| GO:0031326 | Regulation of cellular biosynthetic process | 209 | 2210 | 3.90x10<sup>-02</sup> | 1.34 |
| GO:0010556 | Regulation of macromolecule biosynthetic process | 209 | 2210 | 3.90x10<sup>-02</sup> | 1.34 |
| GO:0019219 | Regulation of nucleobase, nucleoside, nucleotide and nucleic acid metabolic process | 207 | 2192 | 4.83x10<sup>-02</sup> | 1.34 |
| *Other* |  |  |  |  |  |
| GO:0034641 | Cellular nitrogen compound metabolic process | 53 | 373 | 2.05x10<sup>-03</sup> | 2.09 |
| GO:0032501 | Multicellular organismal process | 139 | 1390 | 4.83x10<sup>-02</sup> | 1.41 |
| **Depleted terms** |  |  |  |  |  |
| GO:0007154 | Cell communication | 14 | 1239 | 5.09x10<sup>-22</sup> | 0.14 |
| GO:0030163 | Protein catabolic process | 13 | 1142 | 2.18x10<sup>-20</sup> | 0.14 |
| GO:0009057 | Macromolecule catabolic process | 33 | 1391 | 5.80x10<sup>-14</sup> | 0.29 |
| GO:0009056 | Catabolic process | 50 | 1552 | 9.29x10<sup>-10</sup> | 0.40 |
| GO:0007165 | Signal transduction | 32 | 1184 | 1.00x10<sup>-09</sup> | 0.33 |

<sup>a</sup>Only terms with FDR < 0.05 are shown

